# Shaping a Collaborative, Sustainable, Accessible, and Reproducible Future for Computational Modeling

**DOI:** 10.1101/2025.06.12.656959

**Authors:** Manuel Guidon, Dustin Kaiser, Elisabetta Iavarone, Sylvain Anderegg, Mads Bisgaard, Cedric Bujard, Antonino Cassarà, Pedro Crespo-Valero, Matus Drobuliak, Alessandro Fasse, Yury Hrytsuk, Fariba Karimi, Odei Maiz, Andrei Neagu, Taylor Newton, Javier Ordonez, Tobias Oetiker, Ignacio Pascual, Julian Querido, Sabine Regel, Melanie Steiner, Werner Van Geit, Katie Zhuang, Andrew Weitz, Nicolas Chavannes, Niels Kuster, Esra Neufeld

## Abstract

The o^2^S^2^PARC platform is an open-source, extensible, and scalable cloud-based platform developed in the context of the U.S. National Institutes of Health SPARC program to support collaborative, sustainable, FAIR (findable, accessible, interoperable, reusable) and reproducible computational modeling and analysis. This publication presents the main features of o^2^S^2^PARC, its underlying approaches and philosophy, innovative aspects of the developed technologies, while also drawing attention to its rapid adoption. The paper showcases a variety of applications and use cases enabled by the platform. These include hybrid electromagnetic-electrophysiology simulations of neural interfaces, personalized brain and spinal cord stimulation planning, *in silico* device safety assessments, the training and application of AI systems (e.g., for model-predictive control and medical image segmentation), hybridized surrogate modeling and multi-objective optimization in high-dimensional parameter spaces, sensitive and unbiased validation of measurement devices, and interactive data analysis as paper supplements.

## 2 Introduction

Over the past 30 years, computer-aided engineering (CAE) tools have been successfully applied to optimize medical devices. In recent years, there has been a growing interest in *in silico* approaches in life sciences and medicine, extending far beyond the scope of traditional CAE, for reasons ranging from ethical considerations (e.g., reducing the need for animal experimentation [1]), to advancements in personalized medicine, where subject-specific modeling guide clinical decisions [2]. *In silico* methodologies enable high throughput parameter exploration and can account for inter-subject variability in diverse target populations, producing result with high information density and under controlled conditions. *In silico* studies frequently combine measurement and simulation data and necessitate integration of multiple specialized software tools for data processing, modeling, and analysis [3].

Recent advancements in realistic *in silico* experimentation, digital twins, high performance and cloud computing technologies, and the need for large, often international research collaborations have created new opportunities and requirements. High-performance computing resources are often needed, but demand fluctuations makes purchasing and managing computing clusters inefficient. Adherence to FAIR (Findable, Accessible, Interoperable, and Reusable) principles [4] is as important for computational modeling as for experimental data [5, 6]. Ensuring reproducibility for computational models is challenging, as they frequently involve a large number of steps, specific software requirements, and variations in runtime environments and operating systems. Providing a reproducible runtime environment and guaranteeing that computational pipelines remain functional over time is crucial to ensuring that findings are reproducible and models are sustainably preserved.

Collaboration between research groups is often impeded by the use of diverse software tools, making it difficult to create integrated pipelines. On the other hand, expecting established, optimized, and entrenched tools to be replaced by more generic, cross-disciplinary frameworks (such as CellML [7]), solely for the sake of integration, is unrealistic. Bridging the gap between users that are modeling experts and others (e.g., research clinicians) is even more challenging, but crucial for translational research and for making the benefits of computational modeling accessible to a broader audience.

Several online platforms have been developed to address these challenges. In genomics, projects like the Galaxy Project [8] and bioinformatics workflow managers [9] facilitate the analysis of “next-generation” DNA sequencing data. In other life sciences applications, software for simulating complex systems, such as multicellular interactions (PhysiCell) [10], is available on platforms like nanoHub.org [11]. Platforms like Open Source Brain [12] and brainlife.io [13] provide models, data, and analysis tools for neuroscience research. Real-time collaboration platforms for data analytics, like DeepNote [14], Google Colab [15, 16], and Databricks [17] offer Python notebook-style interfaces but require significant programming experience, which can hinder collaboration with domain experts in life sciences and medicine, as recognized by the developers of the Texera platform [18].

The SPARC program (Simulating Peripheral Activity to Relieve Conditions, [19]), funded by the US National Institutes of Health (NIH) seeks to compensate for lost bodily functionality and advance autonomic nervous system modulation approaches for the treatment of diseases and conditions by unveiling nerve-organ interactions. The SPARC Data and Resource Center (DRC) was established to publish SPARC outcomes through a multifunctional online hub – the SPARC Portal (https://sparc.science). The DRC integrates data and knowledge management, data mapping, and computational modeling. To ensure that computational models and data analyses developed by the >100 SPARC funded research teams adhere to FAIR principles, are accessible to non-modeling-experts, reproducible, and sustainable, a dedicated platform was developed: o^2^S^2^PARC – Open Online Simulations for SPARC. o^2^S^2^PARC serves to establish, host, integrate, execute, share, and publish computational models and data analyses. The platform has since evolved beyond its initial SPARC community and the wider computational life sciences field, enabling numerous and diverse applications across a wide range of disciplines (see Sect. 3.2).

## 3 Results

### 3.1 o^2^S^2^PARC: A Modular, Flexible, and User-Friendly Online Platform

The full functionality of o^2^S^2^PARC and its scalable computational resources are accessible online (https://osparc.io), requiring only a web-browser. Its open-source code (permissive MIT license) can be found on GitHub [20]. Upon logging in, users can access “Studies”, which include their own projects and those shared with them; “Templates”, which are pre-configured Studies shared by others that can be cloned and adapted; and “Services”, which are applications, scripts, data-importers, etc. that serve as the building blocks of “Studies”. Studies, Templates, and Services can be private, shared with one or multiple users, or with an entire “Organization”. Each Service is a reusable module that can function independently (such as a segmentation tool or JupyterLab) or be connected to other Services to build pipelines. The connections between Services are established via their input/output ports, which are specified through standardized metadata (JSON schemas extended with custom properties, such as ‘x-unit’ for units) for input validation, output guarantee definition, and port compatibility checking.

Since its public release in early 2023, o^2^S^2^PARC has seen steadily growing adoption within the computational modeling community. As of March 2025, the platform has surpassed 1,800 registered users, features over 80 published and reusable Services (with many more maintained privately), sees an average of several thousand Service accesses per month, and has generated more than 100 TB of data through o^2^S^2^PARC-hosted pipelines (Fig. 1). Explorable o^2^S^2^PARC Studies have been included as supplements in high impact papers (see Applications), and a range of o^2^S^2^PARC-based free and commercial 3rd party applications have been developed (see Applications and Discussion).

**Figure 1.**
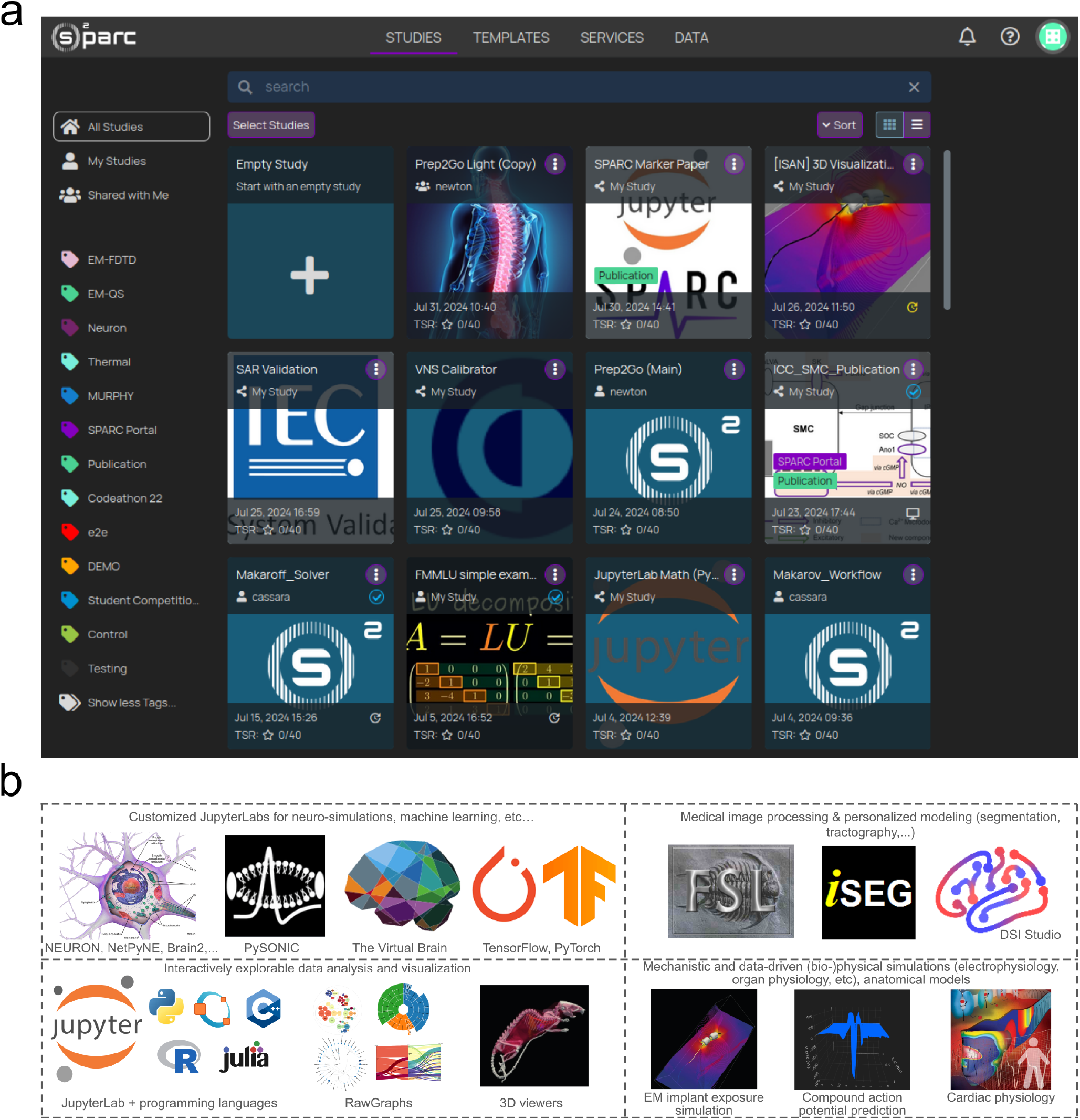
o^2^S^2^PARC Offers a Large Variety of Services for an Increasing Number of Users. **a)** Example of an o^2^S^2^PARC user’s Studies in the Dashboard. **b)** A subset of software, libraries, frameworks and models available as Services on o^2^S^2^PARC. More than 80 such services have been published to date.

Services are categorized into two types: interactive and computational. The former offer a graphical user interface (GUI) and support interactions during run-time. They are used for tasks such as manual segmentation of medical images and inspection of simulation results. The latter execute a predefined task (e.g., a script or solver), making them ideal for automated pipelines and scalable processing, e.g., in the context of meta-modeling (see below). The available Services cover a wide variety of tools commonly used in different life sciences domains: a variety of JupyterLab flavors (e.g., scripting in Python/C++/Julia, Octave, computational neurosciences, artificial intelligence (AI) training and application), medical image processing and personalized modeling (e.g., segmentation, tractography), mechanistic and data-driven (bio-)physical simulation (e.g., electrophysiology, organ physiology), anatomical models, brain network modeling with graph theoretical analysis, and much more (Fig. 1). Service contributors are not forced to open-source their codes, and it is even possible to offer licensed commercial Services.

Besides accessing o^2^S^2^PARC through its web-GUI, its computational resources and Services can be leveraged through a RESTful API for which a dedicated Python clident is available [21], e.g., for application development or high throughput analyses. The platform is designed to suit average users, advanced power-users comfortable with coding, and application developers. Complex, expert-designed pipelines, e.g., for treatment planning, can be transformed into “Guided Applications”, which provide lay users with an intuitive, step-by-step workflow that hides the underlying complexity and only exposes the required interactions.

The platform further includes an extensive quality assurance framework, an associated computational minimal information standard (cMIS; part of SDS 3.0 [22]) and support for the Ten Simple Rules for Evaluating Model Credibility (TSR; [23, 24]), functionality to facilitate collaboration and communication across large consortia, DevOps infrastructure, meta-modeling functionality (e.g., surrogate modeling, uncertainty propagation, sensitivity analysis, multi-goal optimization; benefiting from parallelization on scalable resources), a coupling and control framework (e.g., for model-predictive control [25, 26]), user feedback channels with issue tracking, and more. Additional information can be found in the Methods Section and the user manual [27].

### 3.2 Application Examples

Published modeling applications from the field of bioelectronic medicine include *in silico* assessment of vagus nerve stimulation safety, design of neural interfaces for improved stimulation selectivity, effectivity, and efficiency, interpretation and maximization of neural signal information content, and more. o^2^S^2^PARC has been applied for personalized treatment planing for neuroprosthetics and brain stimulation.

#### 3.2.1. Treatment Planning: Re-usable and Integrated Modeling Pipelines for Personalized Spinal Cord Stimulation (SCS)

Over 15 million people worldwide live with spinal cord injuries (SCI; [28]), resulting in significant disabilities such as partial or complete loss of sensory and motor functions, bowel, bladder, and sexual dysfunction, and dysregulation of blood pressure, heart rate, and body temperature. Epidural electrical stimulation (EES) has shown promise in rehabilitating motor-impaired patients, enabling study participants with SCI to stand, walk, cycle, swim, and control trunk movements again. A sophisticated modeling pipeline involving hybrid EM-electrophysiological simulations was used for pre-operative planning (implant selection, neurosurgical guidance, rapid stimulation configuration of optimized stimulation programs) [29]. Similar modeling was used to design an implant with superior selectivity and effectivity [29, 30], and for interaction mechanism elucidation [31, 32]. Due to vast anatomic and physiologic variability, personalization of these treatments is essential. This is achieved by using MRI data for anatomical model generation (fully automatized, deep-learning-based segmentation), heterogeneous and anisotropic neural tissue properties assignment, fiber tracking, and target identification. Coupled EM and neuronal dynamics simulations subsequently serve to predict spinal root activation selectivity and optimize treatment safety and effectivity. Hybrid surrogate model generation and genetic multi-goal optimization using o^2^S^2^PARC’s meta-modeling framework allows to explore a high dimensional stimulation parameter space [33]. The entire pipeline runs on o^2^S^2^PARC and consists of 25 different Services (Fig. 2).

**Figure 2.**
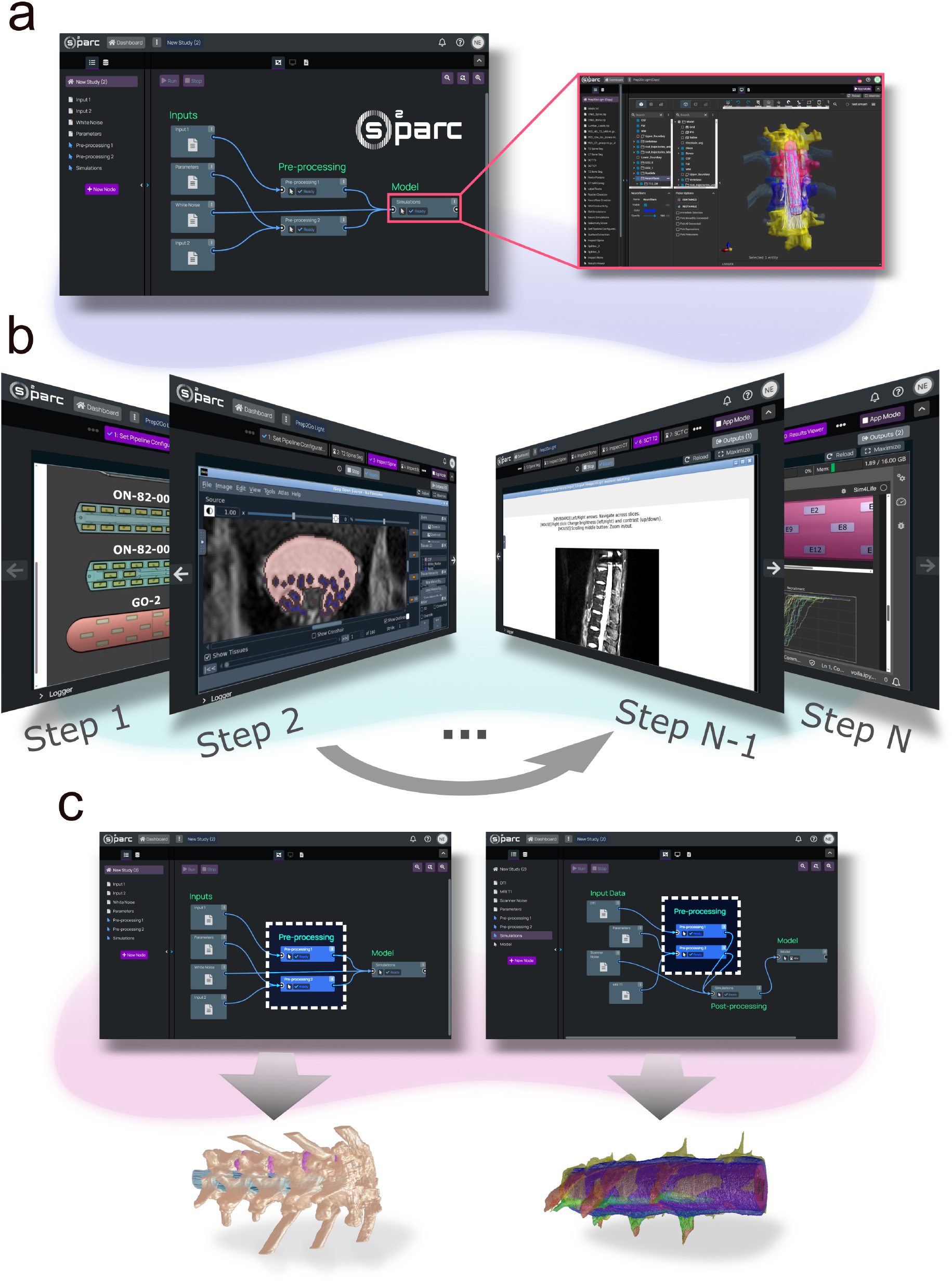
Complex Pipelines and “Guided Applications” for Treatment Planning. **a)** A detailed pipeline composed of 25 coupled Services for spinal cord stimulation (SCS) modeling. **b)** The same pipeline in “Guided Application” mode, exposing only the eight inspection and interaction steps requested by clinical users. **b)** The pipeline designed for SCS stimulation in human to restore locomotion was readily adapted and modules reused for the modeling of feline SCS for bladder control research.

**Figure 3.**
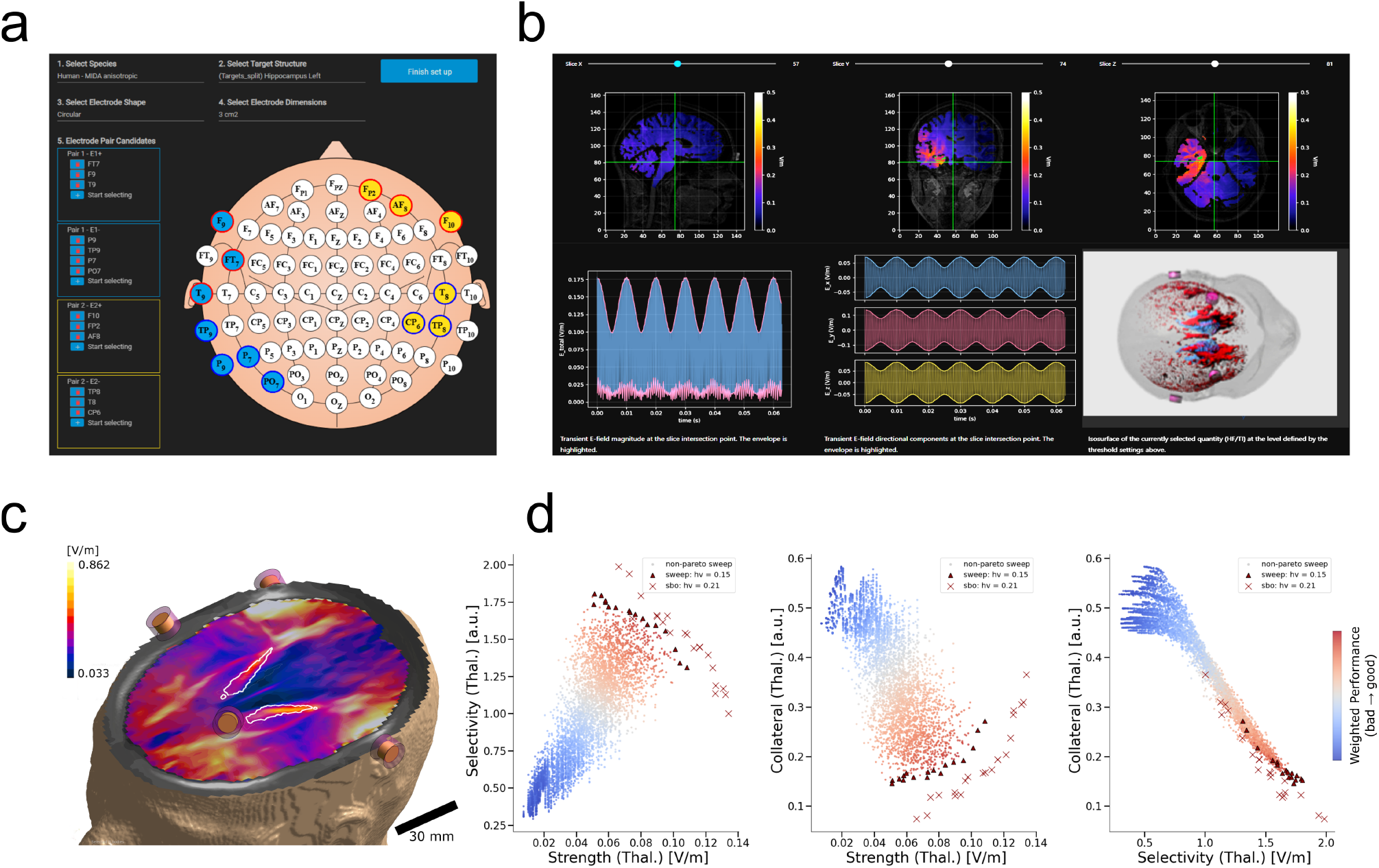
Interactive Workflow for Temporal Interference Planning (TIP) **a)** Setup definition. **b)** Exploration step (partial view only) in which optimized configuration from the multi-goal optimization algorithm can be further refined; the user can interactively weight conflicting goals and adapt stimulation parameters, while visualizing and quantifying their impact on a range of quality metrics. **c)** Cross-section view of an anatomical model and induced field distribution (single channel field). **d)** Scatter plot displaying the performance of investigated configurations in terms of stimulation selectivity, strength, and unwanted off-target stimulation, along with the identified front of Pareto-optimal solutions (triangles: configurations identified using brute force sweeping on limited set of candidate electrodes; crosses: configurations obtained using iterative surrogate model refinement and multi-goal optimization; the latter clearly achieves superior performances and also identifies a broader front of solution – with the added benefit of being faster, systematic, unbiased, and suited for inexperienced users).

To support research clinicians, a “Guided Application” mode was added. While segmentation, simulation setup, solving, optimization, and analysis are computational and executed in the background, eight interactive steps are exposed, providing implant selection options, segmentation and model generation inspection, the possibility of manually adapting segmentation and implant placement (will be removed, once confidence is established), interactive weighting of conflicting optimization goals, inspection of fields, fiber recruitment, and treatment quality metrics for the best rated configurations, and report generation. Usability testing involved three clinicians familiar with the intervention, but never before exposed to the modeling pipeline. They received a one page primer and were all able to successfully complete the planning.

Another o^2^S^2^PARC-based pipeline showcases one of the platform’s core principles: re-usability. Aspects of the pipeline developed for locomotion restoration were reused in an animal study on SCS for bladder control in cats [34, 35], which retained certain Services (e.g., EM-neuron interaction modeling), adapted others (e.g., AI segmentation), and introduced new ones, e.g., to leverage diffusion tensor imaging (DTI) data for fiber tracking. This illustrates how o^2^S^2^PARC supports FAIR principles, benefiting diverse medical applications.

#### 3.2.2 Guided Applications for Non-Invasive Brain Stimulation

Deep brain stimulation (DBS) techniques using implanted electrodes are frequently employed to treat severe neurological and psychiatric disorders. However, their invasiveness and the associated risks limits wider clinical adoption. Temporal Interference (TI) is an emerging non-invasive stimulation technique that uses electric fields at kHz-frequencies too high to effectively neuromodulate on their own, but with frequency offsets and/or phase modulation in the physiological range, which results in amplitude modulation that neurons respond to. By combining multiple current pathways, their overlap region where modulation arises can be preferentially targeted to deep brain regions without affecting overlying structures [36]. Despite uncertainty about its mechanistic basis, TI has attracted significant interest in performing *in vitro*, animal, and human studies, creating a demand for computational tools to optimize stimulation protocols and elucidate TI mechanisms.

The o^2^S^2^PARC-powered, online-accessible Temporal Interference Planning (TIP) tool facilitates the quantification and optimization of *in vivo* TI exposure through a three step “Guided Application” (Fig. 2), i.e., (1) definition of target brain region(s) and constraints on potential electrode placement; (2) identification of an optimized stimulation protocol through advanced EM simulations using detailed personalized (anatomy and heterogeneous, anisotropic brain property maps derived from MRI data) or reference head models (e.g., MIDA [37] with tissue properties from [38]; multiple reference models capture inter-subject and -species variability); (3) interactive in-depth exploration (e.g., dosimetric quantities of hypothesized mechanistic relevance in different atlas-based brain regions). Step (2) employs multi-goal optimization considering target exposure magnitude, stimulation selectivity, and minimization of undesired collateral stimulation, and empowers users to interactively weigh conflicting goals, while visualizing TI and high-frequency fields and quality metrics. In addition, a report with quantitative and visual results on the preferred conditions can be generated. Recent releases also introduced support for multichannel TI (up to 8 channels for enhanced focality, selectivity, and multifocal stimulation, distributing currents over larger scalp areas to minimize confounding sensations) and phase-modulation TI (for enhanced stimulation effectivity and to selectively activate specific neural subpopulations).

#### 3.2.3 Explorable Data Analyses for Reproducible and FAIR Research

Exploration and publication of experimental and/or simulation data often requires extensive preand post-processing, such as statistical analysis, visualizations, and uncertainty quantification, which is typically performed using custom software. While code repositories such as GitHub and Zenodo are available [39], they do not provide an easy way for researchers to reproduce results, or perform additional, in-depth investigations. o^2^S^2^PARC permits to supplement publications with explorable results and analyses. Templates can be openly published, resulting in a unique reference link that can be included in a paper. The content can range from pre-configured model and result viewers, to publishing the code used for analysis and figure generation, or entire modeling pipelines. Code publication can take the form of online accessible JupyterLabs (similar to Google Colab [28]) that are either fully open, providing researchers with the flexibility to adapt or reuse published code, or in a preconfigured and more constrained, but still interactive Voilà mode [40]. For example, [29] has an o^2^S^2^PARC Supplement that allows interactive inspection of spinal cord models from 15 healthy volunteers with overlaid EM simulation results, which can be used to study the influence of electrode positioning on the recruitment of dorsal spinal cord roots. A recent publication on TI stimulation of the human hippocampus [41] uses o^2^S^2^PARC to allow viewing of head models with distinct brain regions and exploration of EM simulation results (spatial distribution of amplitude modulation magnitude and carrier fields strength) optimized to preferentially target different hippocampal sub-regions, as confirmed experimentally.

The SPARC Portal hosts a number of computational models [42] that can be executed on o^2^S^2^PARC or even directly on the SPARC Portal through o^2^S^2^PARC’s computational API. For example, a recent publication features a model of slow waves and phasic contractions in the distal stomach ([43], "https://doi.org/10.26275/ixhd-0pfa" DOI:10.26275/ixhd-0pfa). Users can set simulation parameters through an intuitive GUI, run the simulations using the OpenCOR [44] Service on o^2^S^2^PARC, and interactively explore output quantities (e.g., evolution of gastric tension over time). Reconfigurable models on the SPARC Portal include JupyterLab Services, where users can modify the code, an extended version of the cardiovascular system model by Haberbusch et al. ([45], "https://doi.org/10.26275/5yvu-tr0d" DOI:10.26275/5yvu-tr0d), “Guided Applications” with interactive coding environment enriched by “Instructions”, and complex simulation pipelines, such as ASCENT (Automated Simulations to Characterize Electrical Nerve Thresholds) for vagus nerve stimulation modeling ([46, 47], "https://doi.org/10.26275/0jz3-zrlo" DOI:10.26275/0jz3-zrlo; Fig. 4). Despite depending on the commercial COMSOL software, ASCENT runs on o^2^S^2^PARC without necessitating the user to install any software or licenses, thanks to a pre-configured license servers on the platform and an agreement that allows running predefined and restricted ASCENT scripts, in which only specific parameters can be modified. However, a series of alternative open-and closed-source solvers is also available on o^2^S^2^PARC.

**Figure 4.**
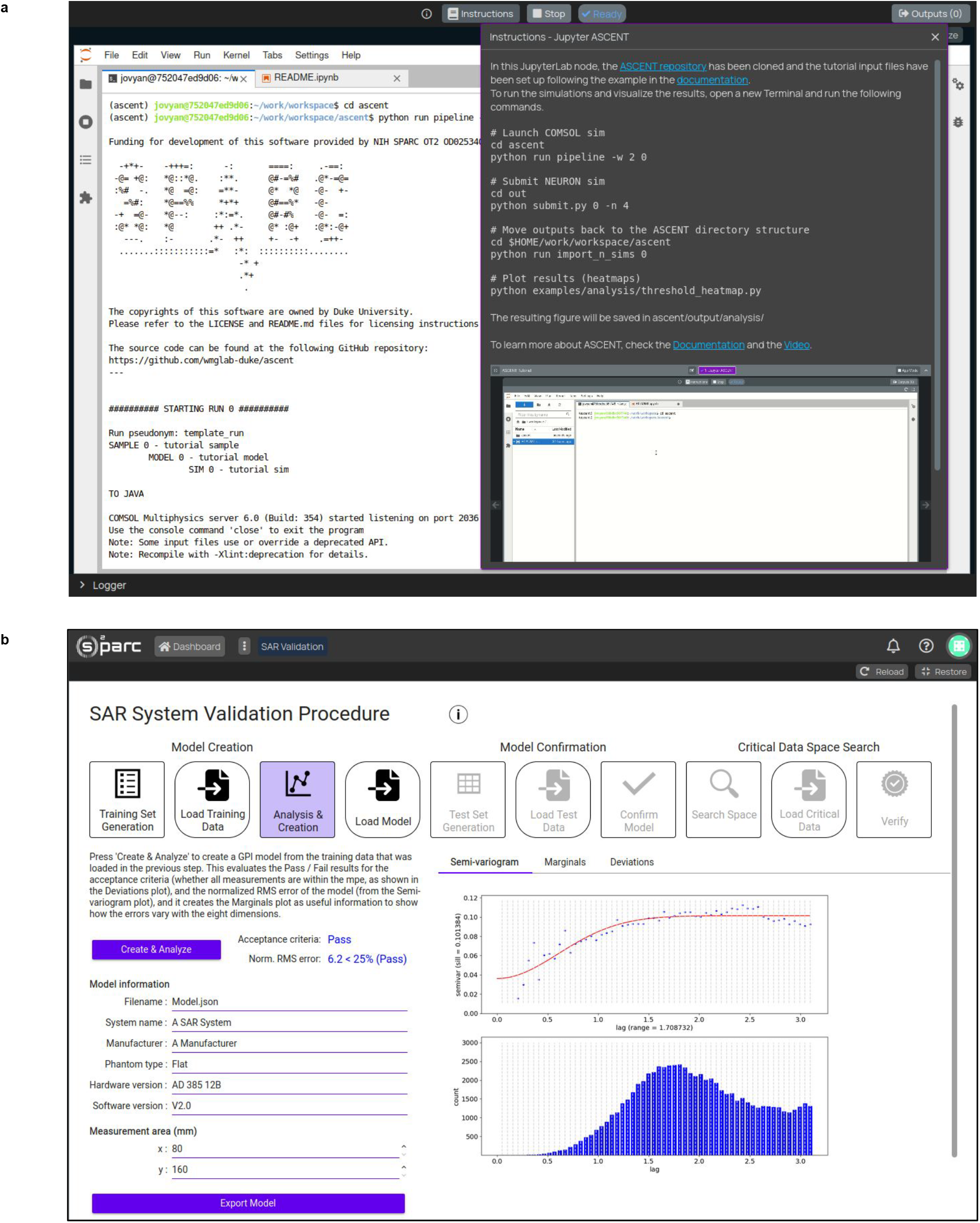
Complex modeling and compliance assessment tools made accessible and user-friendly on o^2^S^2^PARC. **a)** ASCENT [46, 47] includes a flexible JupyterLab environment featuring an “Instructions” GUI that supports the display of rich text, images, animations and short movies. **b)** Customized GUI for efficient, comprehensive, and technology-agnostic validation of specific absorption rate (SAR) measurement systems, proposed to the IEC to address gaps in current standards [48, 49]. Guided model creation, measurements collections, modeling testing a validation is accompanied by detailed analyses and export of detailed reports.

#### 3.2.4 Device Validation Using Gaussian Process Modeling

o^2^S^2^PARC has also been utilized to develop self-contained GUI applications unrelated to life sciences. An example is https://sarvalidation.site, a reference implementation of a novel method designed to maximize the likelihood of detecting measurement errors exceeding a given tolerance threshold [50]. This method is likely to be adopted during revisions of the standards defining the exposure limits and the requirements for specific absorption rate (SAR) measurement systems used in compliance testing for wireless devices [48, 49]. Since different measurement technologies will be permitted, the validation procedure must be unbiased, systematic, and technology-agnostic. It leverages Gaussian Processes, a form of surrogate modeling, to manage the high-dimensional parameter spaces associated with complex SAR measurements. The mathematical formulation of this approach is intricate and a user-friendly reference implementation was required to allow test labs, as well as system and device manufacturers, to assess its practicality. The SAR validation tool, with its intuitive GUI, guides users through the specified three steps: (1) Model creation, which requires users to specify the frequency range, measurement area, and sample size, thereby generating a set of test conditions. Once the conditions are defined and actual measurement data are collected with the device, users can create the model and evaluate the pass/fail results against the acceptance criteria. (2) Validation of the model with a new set of measurements and tests (i.e., data not used to generate the model) to confirm its accuracy. (3) Identification of the most critical regions of the test space, where measurements are more likely to exceed the maximum permissible error, as defined in IEC 62209-3 [48]. The o^2^S^2^PARC application (Fig. 4) includes interactive data visualization, is fully automated and allows users to generate and download PDF reports, e.g., for regulatory approval.

## 4 Discussion

o^2^S^2^PARC is designed to address fundamental challenges in computational modeling across the life sciences and beyond, offering an integrated, user-friendly, and powerful resource to a growing user base. By facilitating collaborative, shareable, sustainable, reproducible, and FAIR research (Tab. 4), o^2^S^2^PARC aims to accelerate the development and dissemination of high-quality computational models and methods, and to allow modelers to leverage and build on top of existing contributions from others.

Quality assurance is integral to o^2^S^2^PARC’s code development and operations. The platform’s code-base boasts a high test coverage (87% as of March ‘25,), ensuring that code changes do not negatively impact existing functionality. Adhering to the latest coding best practices, such as type checking and code formatting, provides an additional layer of automatic quality assurance, minimizing human errors and enhancing the effectiveness of code reviews. Additionally, automated tools perform code quality and security analysis on each code change, with these metrics being openly available on the GitHub repository through “badges” [20]. The overall platform is also rigorously tested via end-to-end tests that simulate typical user workflows, ensuring stability and reliability for end users. An established release workflow, including staging and production deployments, further ensures a smooth and stable user experience, with high automation minimizing human errors and reducing the time for code changes to reach production. Various channels for user feedback result in git-tracked bug reports and feature requests.

o^2^S^2^PARC closely integrates with the SPARC Portal. Datasets from the Portal can directly be imported, computational models published on the Portal can be executed on o^2^S^2^PARC, o^2^S^2^PARC Services provide access to SPARC anatomical and functional maps, and the Portal leverages o^2^S^2^PARC to launch small, interactive simulations and display results directly on the Portal.

o^2^S^2^PARC (`osparc.io`) is open-sourced under the permissive MIT license, leverages open-source tools, and facilitates the integration of open-source software as Services. Projects developed on the platform can be published, promoting open science. Free and commercial platforms based on o^2^S^2^PARC’s code-base and technologies have been deployed, such as a stand-alone TIP platform, and different variant of the Sim4Life computational life sciences software (ZMT Zurich MedTech AG, Switzerland [3]). S4L.lite has reduced functionality, but is free for students (AWS resource costs are sponsored), S4L.sciences for academic users attributes resource costs to users, but does not charge license fees, while pay-per-use Sim4Life.web brings the entire functionality of the desktop Sim4Life software to a scalable, web-accessible cloud resource and allows advanced multiphysics simulations in complex environments, such as the human body. Such commercial applications are fundamental to sustainably support continued o^2^S^2^PARC development and access to published models beyond NIH SPARC program support.

A key o^2^S^2^PARC intent is to make the user experience of conducting computational science superior to the traditional one of having to set up coding environments and tools, of manually integrating different steps, of setting up code repositories for collaboration, and of managing jobs on a high-performance computing cluster. There is an intrinsic value perceived by the users in being able to access their work from any device that contains a web-browser, without the need to set up specific tooling like Virtual Private Network tunnels or install specialized software on every computer they work with. Shifting the provisioning and choice of servers away from users enables them furthermore to focus their attention on their core tasks, eliminating worries about hardware compatibility, licensing servers, or software updates. However, this promise of a better user experience can only manifest if software such as o^2^S^2^PARC is able to ensure consistent availability, reliability, and stability, and if researchers can continue to use the software they are accustomed to. Only then will o^2^S^2^PARC have a chance of replacing custom setups, which explains the current development focus on usability, platform hardening, and sustainability. In parallel, the growing user-base is leveraged to obtain feedback that helps improve the platform experience, usefulness, and operation.

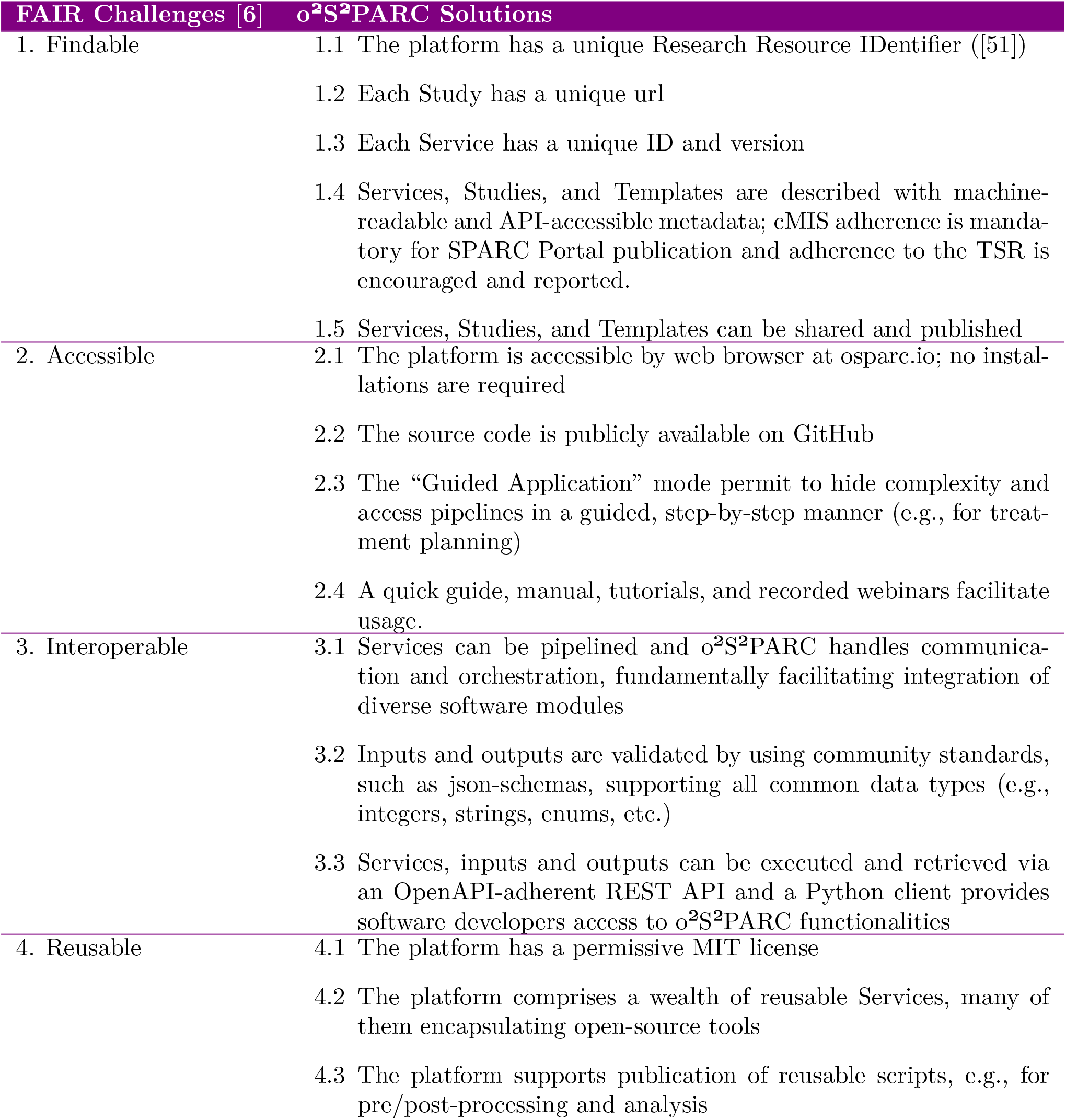
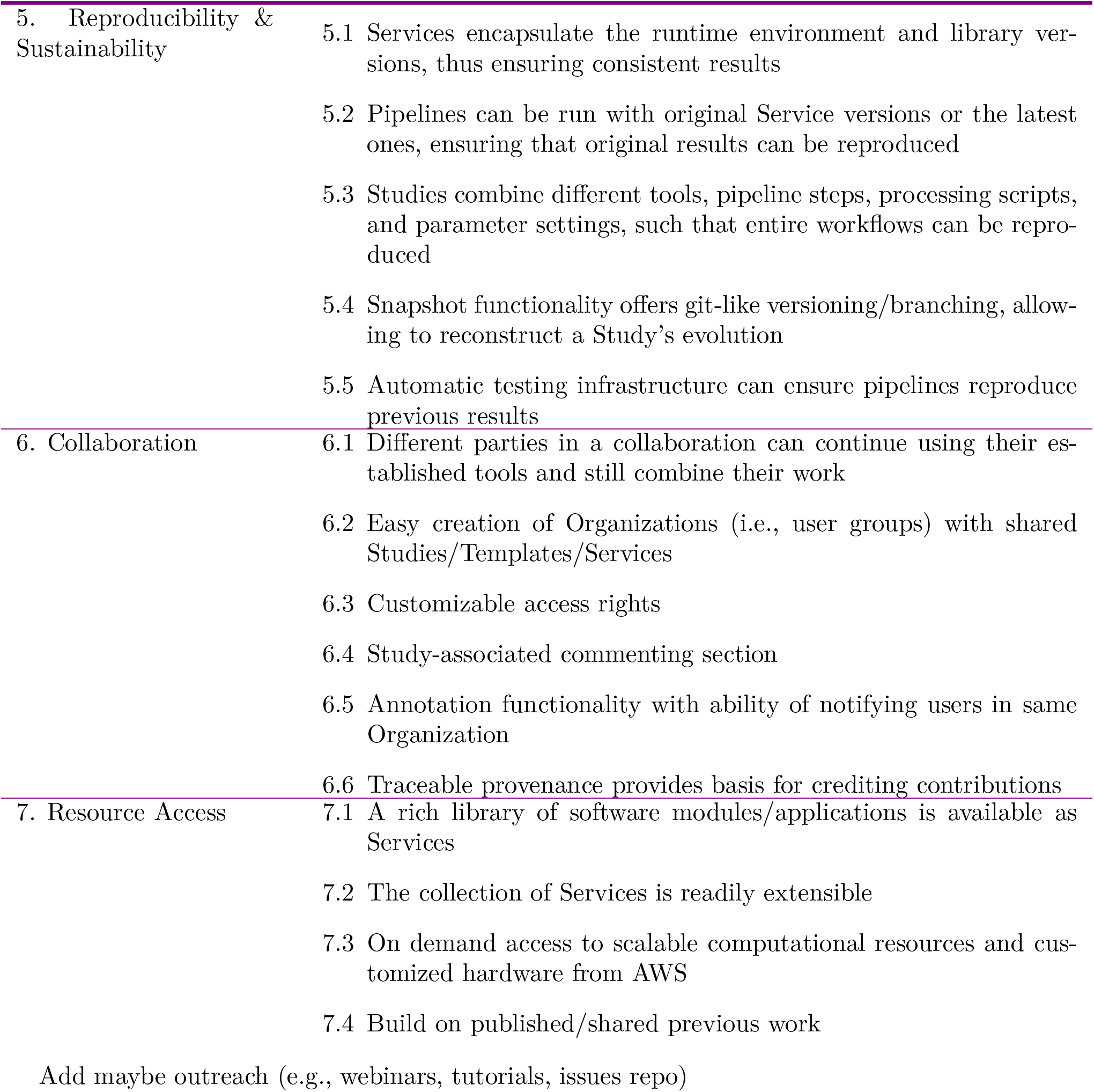

| FAIR Challenges [6] | o<sup>2</sup>S<sup>2</sup>PARC Solutions |
| --- | --- |
| 1. Findable | 1.1 The platform has a unique Research Resource IDentifier ([51])<br>1.2 Each Study has a unique url<br>1.3 Each Service has a unique ID and version<br>1.4 Services, Studies, and Templates are described with machine-readable and API-accessible metadata; cMIS adherence is mandatory for SPARC Portal publication and adherence to the TSR is encouraged and reported.<br>1.5 Services, Studies, and Templates can be shared and published |
| 2. Accessible | 2.1 The platform is accessible by web browser at osparc.io; no installations are required<br>2.2 The source code is publicly available on GitHub<br>2.3 The “Guided Application” mode permit to hide complexity and access pipelines in a guided, step-by-step manner (e.g., for treatment planning)<br>2.4 A quick guide, manual, tutorials, and recorded webinars facilitate usage. |
| 3. Interoperable | 3.1 Services can be pipelined and o<sup>2</sup>S<sup>2</sup>PARC handles communication and orchestration, fundamentally facilitating integration of diverse software modules<br>3.2 Inputs and outputs are validated by using community standards, such as json-schemas, supporting all common data types (e.g., integers, strings, enums, etc.)<br>3.3 Services, inputs and outputs can be executed and retrieved via an OpenAPI-adherent REST API and a Python client provides software developers access to o<sup>2</sup>S<sup>2</sup>PARC functionalities |
| 4. Reusable | 4.1 The platform has a permissive MIT license<br>4.2 The platform comprises a wealth of reusable Services, many of them encapsulating open-source tools<br>4.3 The platform supports publication of reusable scripts, e.g., for pre/post-processing and analysis |

| CLS Challenges | o<sup>2</sup>S<sup>2</sup>PARC Solutions |
| --- | --- |
| 5. Reproducibility & Sustainability | 5.1 Services encapsulate the runtime environment and library versions, thus ensuring consistent results<br>5.2 Pipelines can be run with original Service versions or the latest ones, ensuring that original results can be reproduced<br>5.3 Studies combine different tools, pipeline steps, processing scripts, and parameter settings, such that entire workflows can be reproduced<br>5.4 Snapshot functionality offers git-like versioning/branching, allowing to reconstruct a Study's evolution<br>5.5 Automatic testing infrastructure can ensure pipelines reproduce previous results |
| 6. Collaboration | 6.1 Different parties in a collaboration can continue using their established tools and still combine their work<br>6.2 Easy creation of Organizations (i.e., user groups) with shared Studies/Templates/Services<br>6.3 Customizable access rights<br>6.4 Study-associated commenting section<br>6.5 Annotation functionality with ability of notifying users in same Organization<br>6.6 Traceable provenance provides basis for crediting contributions |
| 7. Resource Access | 7.1 A rich library of software modules/applications is available as Services<br>7.2 The collection of Services is readily extensible<br>7.3 On demand access to scalable computational resources and customized hardware from AWS<br>7.4 Build on published/shared previous work |

Add maybe outreach (e.g., webinars, tutorials, issues repo)

## 5. Methods

### 5.1 Code design and implementation

o^2^S^2^PARC’s code consists of a frontend, a backend, servers, processes that manage, e.g., the execution of jobs and data storage, and a “Cluster Monitoring Framework”, which provides observability and metrics on the state, usage, and health of the platform. The following is a high-level overview; further information can be found in the source-code repository (https://github.com/topics/osparc) and its associated documentation.

### 5.1 Frontend

By frontend we understand the web-page that a user interacts with. This JavaScript code is executed inside the user’s browser (client-side code) and permits visualization of and interaction with content, such as data, scientific results, or information about computational pipelines. Content is stored on and retrieved from remote servers (“backend”). Frontend-backend communication employs REST APIs according to CRUD (Create, Read, Update and Delete) principles [52], and bi-directional realtime communication, e.g., for logs and state changes initiated by the backend, is facilitated using web-sockets. As javascript framework, qooxdoo was chosen [53], which stands out through its focus on object-oriented design and fully non-commercial, independent nature.

#### 5.1.2 Backend

The o^2^S^2^PARC backend is best described as a multi-tenant microservice web-application built mainly in Python. It is a suite of applications, composed of a multitude of small, autonomous sub-programs that communicate using standardized and versioned APIs. Microservice architectures were introduced to avoid shortcomings of monolithic software [54]. A main benefit, particularly for multi-tenant web-applications, is that microservices facilitate scaling with increased usage/load (horizontal scaling) through addition of additional instances and servers. o^2^S^2^PARC’s microservices run as containers on linux hosts. Containerization adds another layer of encapsulation, abstraction and security, but is considerably more robust than secluding the code into, e.g., virtual machine [55, 56].

Figure 5 illustrates the o^2^S^2^PARC backend design and interactions with/between its microservices. Key o^2^S^2^PARC microservices include: a *webserver*, which is a group of services with different responsibilities such as serving the webpage to the user and exposing a dedicated API (web-api) for it; a *director* that orchestrates the execution of Services and computational jobs; a *storage* service, which handles placement and retrieval of uploaded or user-generated data; an *api-server* that exposes a separate public-api alternative to the web-api for programmatic access to o^2^S^2^PARC content and functionalities; as well as a *catalog* storing and caching the available Services.

**Figure 5.**
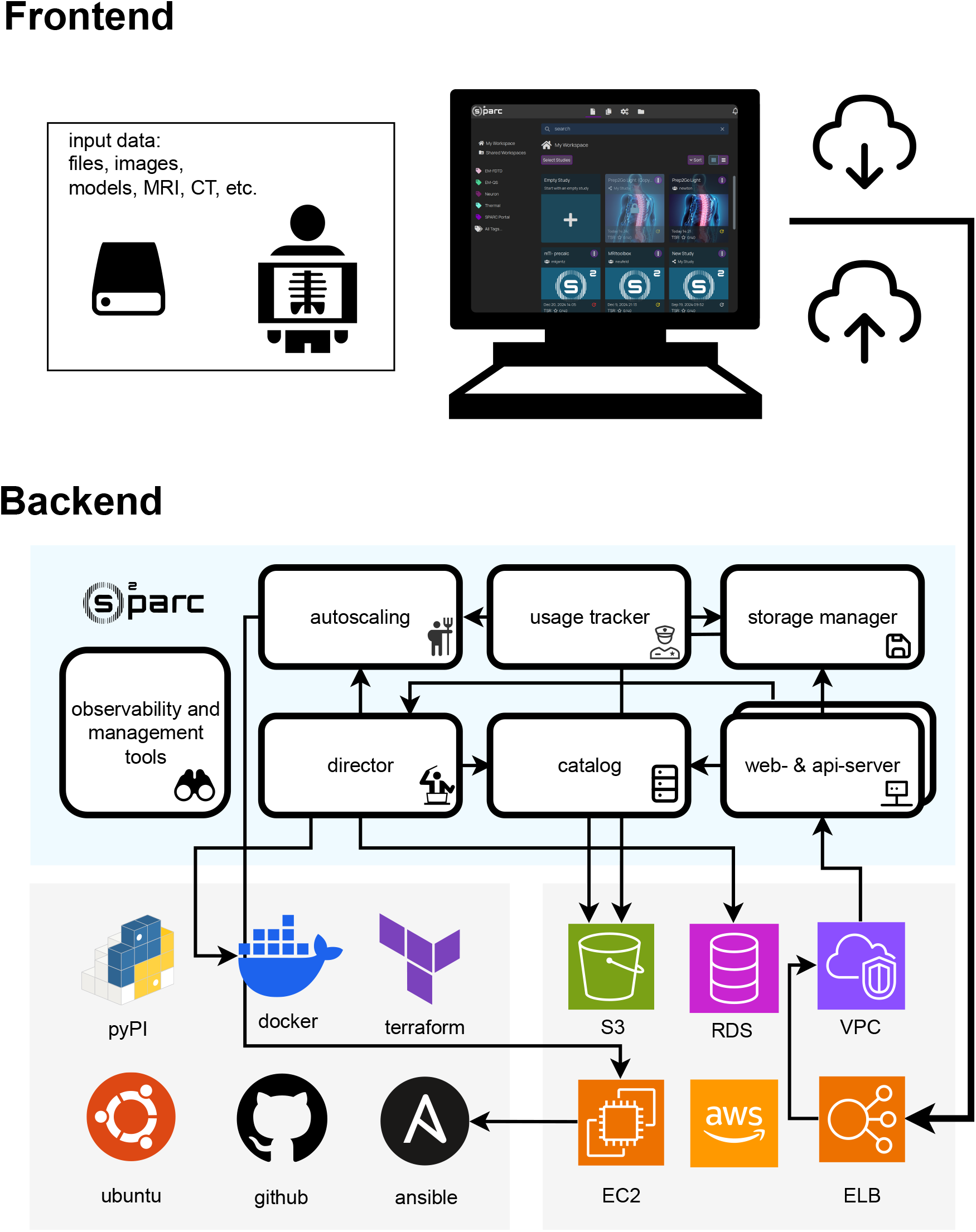
Schematic overview of the o^2^S^2^PARC code architecture. Highlighting the microservice design of o^2^S^2^PARC (blue panel), leveraged AWS cloud services, as well as third-party dependencies and integrations. Arrows representing possible flow of information via remote APIs. Not all o^2^S^2^PARC microservices are shown. For more detailed information on the software, see text. S3: Simple Storage Service, RDS: Relational Da1ta5base Service, VPC: Virtual Private Cloud, ELB: Elastic Load-balancing, EC2: Elastic Compute Cloud; ubuntu: GNU/linux operating system, pyPI: Python Package Index, terraform: infrastructure-as-code software, ansible: configuration management software, github: distributed version control platform, docker: container virtualization software.

#### 5.1.3 Management stack

Software-as-a-Service platforms such as o^2^S^2^PARC are known to degrade, and with growing complexity the number of corner-cases and unexpected occurrences increases rapidly [57]. Thus, it is crucial to monitor state, health, and usage. While o^2^S^2^PARC *can* be run in a stripped bare-bones version, the servers of osparc.io run a plethora of additional tooling to support monitoring and if necessary intervention, and to ensure server and data security.

o^2^S^2^PARC stores its state in two databases: a redis key-value store [58] and a relational postgres database [59]. Further information is stored in the state of the container orchestration engine, in this case docker. Browser-based tools enable devops to inspect, dump, or manipulate these stores to perform ad-hoc maintenance in a surgical manner. Graylog [60] is used for log-management tool and search. Usage and performance metrics are continuously exported from o^2^S^2^PARC microservices into a specialized time-series database (prometheus [61]) and can be aggregated and visualized in grafana [62] dashboards.

For security reasons, all interactions with the software happen by means of a reverse-proxy (traefik [63]), a tool that distributes and inspects incoming web requests. It handles access rights, load-balancing of incoming requests, and rate-limiting in case of request-floods or denial-of-service attacks. For the operation of o^2^S^2^PARC, a heavily customized configuration of the reverse-proxy is necessary to meet the desired security and availability levels.

To identify nascent security issues or vulnerabilities in the code or its third-party dependencies, it is crucial to have full control over the precise version of each software-dependency running at any time. This is achieved by employing the ansible automation software [64] to implement the infrastructure-as-code paradigm [65]. The idempotent automation code developed for o^2^S^2^PARC does not affect services that are already in their desired state, such that it can be run at any point in time, not only to provision new, unconfigured servers, but also to restore the desired state of any computer. Automatic tooling parses server states to detect security issues and raise alerts. For cloud deployments, such as osparc.io on AWS, the virtual cloud architecture layout (virtual private clouds, subnets, network interfaces, machines, route tables, etc.) is similarly compiled in code (terraform [66]).

The o^2^S^2^PARC back-end code relies extensively on unit-, integration-and end-to-end testing to detect potential failures early in the development lifecycle, before breaking code changes are published to the production environment, where users interact with the application. End-to-end tests simulate actual user interactions with the platform programmatically on an hourly basis.

Continuous deployment [67] pipelines, which roll out desired changes to the remote servers in a fully predictable and automated fashion, also help ensure smooth operation. Any code change is rolled out without the need for human intervention on o^2^S^2^PARC’s development deployment, thus allowing developers to see their code in action within an hour after committing their changes to the source-code repository. Releases to the production-servers that run osparc.io use identical workflows, though they are triggered manually after prior announcement.

### 5.2 Key Technological Design Aspects

A series of innovative key technologies are fundamental to the platform:

#### 5.2.1 Core Platform Architecture

The o^2^S^2^PARC backend is a distributed web application built on a microservice architecture, primarily developed in Python. It consists of small, autonomous services that communicate via standardized APIs, enabling horizontal scaling and efficient handling of dynamic workloads. Each microservice runs as a Docker container on Linux hosts, leveraging containerization for abstraction, security, and performance. The platform employs Docker Swarm (in some instances also Kubernetes) for container orchestration, ensuring seamless deployment, efficient resource allocation, and dynamic scaling. Key services include a web server for relaying user interaction, a director for orchestrating computational and interactive Services, a storage service for data management, an API server for programmatic access, and a catalog for service discovery. These components communicate via message queues or OpenAPI-compliant transport protocols [68]. Relying on API communication within the o^2^S^2^PARC code ((see Figure 5)) allows the platform to easily interact with third-party services and software, facilitating to use of existing solutions wherever possible (e.g., AWS S3 REST API communication with cloud storage; Pennsive [69] API to access SPARC data).

#### 5.2.2. Pipelining

o^2^S^2^PARC enables the creation and execution of computational pipelines by connecting Services into workflows. Users can design pipelines via an intuitive interface, automating data processing and simulations. The director service orchestrates these workflows, ensuring proper execution order and data flow (files and database tables). Real-time updates, including logs and error messages, are relayed to the user interface, providing transparency and insight into pipeline execution.

#### 5.2.3. Heterogeneous Service Hosting

The platform supports a diverse range of Services with varying resource needs, from lightweight computations to resource-intensive tasks requiring parallel processing (OpenMP, MPI), GPU acceleration (CUDA), OpenGL/Vulkan rendering, and high-bandwidth video streaming. This flexibility allows o^2^S^2^PARC to cater to a broad spectrum of scientific and technical applications. Consequently, the platform must accommodate a wide variety of hardware configurations to meet these diverse requirements effectively.

#### 5.2.4. Dynamic Resource Allocation with Smart Autoscaling

A smart autoscaling mechanism dynamically manages resources, while allowing users to choose from a default setup or customized AWS EC2 instance types. The system is cloud-agnostic, supporting future integration with other providers. Compute nodes are automatically added during high load, maintaining performance. The platform leverages Dask as a scheduler for job queues, dynamically creating temporary Docker Swarm clusters via the autoscaling microservice to efficiently scale workloads. The autoscaling service interacts with AWS’s EC2 API to create and destroy virtual servers on demand. (see Figure 5).

#### 5.2.5 Advanced Data Management

Data-intensive applications are supported through a multi-stage caching system. Transferring large datasets from AWS S3 to compute nodes and synchronizing back can be time-consuming, especially when dealing with terabytes of data. To address this, the platform uses a distributed warm filesystem (AWS EFS) to cache frequently accessed data, enabling fast, mountable access directly on EC2 instances. Similarly, Docker images are pre-pulled onto AWS EBS volumes and attached to compute nodes when needed, reducing startup times and improving efficiency.

#### 5.2.6 Resource Tracking

The platform monitors user consumption of compute, data, and software resources, enabling precise cost tracking and attribution of resource costs (currently covered by the NIH STRIDES Initiative) and the eventual licensing cost of commercial Services to individual users or groups. This information is collected from several microservices across the platform, centralized and managed by the resource tracking service. For services that require significant resources, an optional billing extension is available. It lets users pay either upfront or per use directly through the platform. Some services/tools follow a partial pay-per-use model, e.g., the TIP tool is free when using precomputed head models, but users who need personalized (MRI-based) exposure planning are charged about 30 USD to cover the cost of *∼*70 high-resolution EM simulations on AWS.

#### 5.2.7 Onboarding of User Services

The o^2^S^2^PARC platform supports seamless extension by allowing new Services to be added to its catalog. Each service must follow a defined protocol, specifying input and output ports along with their types (e.g., integers, floats, files). Interactive Services may require additional configurations, such as specifying interconnected containers. In its final form, a Service must be deployable as a Docker container. To streamline onboarding, o^2^S^2^PARC offers cookiecutter templates, simplifying the development and deployment of Services. Once a user Service is published in the catalog, the owner can share it with other users as detailed above.

## 6 Acknowledgments

This work was supported by the National Institutes of Health (NIH) through award OT3OD025348.

## 7 Disclaimer

Andrew Weitz contributed to this work when he was an NIH employee; however, he is no longer affiliated with the NIH.

